# Recycling Materials for Sustainable DNA Origami Manufacturing

**DOI:** 10.1101/2024.06.05.596853

**Authors:** Michael J. Neuhoff, Yuchen Wang, Nicholas J. Vantangoli, Michael G. Poirier, Carlos E. Castro, Wolfgang G. Pfeifer

## Abstract

DNA origami nanotechnology has great potential in multiple fields including biomedical, biophysical, and nanofabrication applications. However, current production pipelines lead to single-use devices incorporating a small fraction of initial reactants, resulting in a wasteful manufacturing process. Here, we introduce two complementary approaches to overcome these limitations by recycling the strand components of DNA origami nanostructures (DONs). We demonstrate reprogramming entire DONs into new devices, re-using scaffold strands. We validate this approach by reprogramming DONs with complex geometries into each other, using their distinct geometries to verify successful scaffold recycling. We reprogram one DON into a dynamic structure and show both pristine and recycled structures display similar properties. Second, we demonstrate the recovery of excess staple strands post-assembly and fold DONs with these recycled strands, showing these structures exhibit the expected geometry and dynamic properties. Finally, we demonstrate the combination of both approaches, successfully fabricating DONs solely from recycled DNA components.

## Introduction

Over the last decades, DNA origami has emerged as a powerful and transformative bottom-up technology to fabricate structures on the nanoscale. (1) This technology harnesses the inherent self-assembly properties of DNA molecules to create nanostructures with remarkable precision and complexity. DNA origami nanostructures (DONs) have demonstrated strong potential in materials science, biophysics, and biomedical applications (2–4) and have been applied to specific tasks such as manufacturing therapeutics, (5–8) realizing templates for optical and electronic materials, (9–11) and creating nanorobots. (12–14) As the applications of DONs continue to expand, there will be an increasing need for larger scale production, leading to a higher demand for DNA raw materials and a concomitant increase in costs and waste associated with the fabrication process. Methods to improve efficiency of material use can play an important role in reducing cost for both early-stage research and for manufacturing scale-up. Furthermore, there is an increasing appreciation for the need to minimize waste in biomaterial manufacturing processes to mitigate potential environmental impacts. (15) Here we demonstrate key steps to advance the sustainable and cost-effective manufacturing of DONs through straightforwardly implementable methods for recycling the constituent DNA components.

One popular framework for synthesizing DNA nanostructures is scaffolded DNA origami where up to 200 short synthetic single-stranded DNA (ssDNA) oligonucleotides, referred to as staple strands, are combined with one long ssDNA component, referred to as a scaffold strand. (16) In this approach, discontinuous base-pairing interactions between the scaffold and staple strands drive folding into a target shape. The fabrication of DONs is performed with a large excess of staple strands, typically 10-fold, to drive the assembly reaction to completion, (1,16,17) meaning around 90% of staple strands are discarded as waste. Typical DON folding reactions in research laboratories yield micrograms of material. (18) While this is sufficient for many basic scientific applications, such as research tools to probe biomolecular interactions, other applications require milligrams or more of DNA, especially to scale to clinical or industry needs, resulting in much larger levels of waste. (18,19) Furthermore, recent work has seen an increase in custom scaffold strands with modifications that include certain sequences as binding domains, sequence motifs to allow downstream processing, or custom sizes to better match specifications of the target application. (6,20–23) Previous studies have estimated the costs of making DONs to be roughly $100-150 per milligram using typical laboratory scale production methods with the major costs being for scaffold and staple strands (18,24). These cost estimates assume in-house production of scaffold strands, which can be tedious and requires specialized equipment. (24,25) Scaffold is available from multiple vendors, but commercial purchase of scaffold more than doubles the production costs of DONs. Hence, methods to effectively recycle both scaffold and staple strands could prove highly useful to maximize use of resources, reduce research and development costs, and mitigate excessive nucleic acid waste.

While recycling of DON components for completely new structures has not previously been reported, several studies have developed designs where structure conformations can be adjusted through actuation mechanisms such as change of pH, temperature, or salt concentration. (26–28) However, these actuation and partial refolding strategies lead to only small changes or local adjustments to the conformation and do not allow complete reprogramming into a new structure with new properties. (26,29–31) Several studies have investigated folding DONs with staple excesses below 10x. (24,32–35) However, the majority of work with DONs uses 10x staple excess to promote folding since the success of lower excess folding is inconsistent and dependent on particular DON design. (24,32,34) Hence, methods to recover and re-use both staple and scaffold strands are important for reliable fabrication while reducing waste. These considerations combine to imply the importance of understanding the impact recycled materials have on DON fabrication and properties, akin to macroscopic material recycling efforts. (36)

Here, we report approaches to recycle both scaffold and staple strand components and demonstrate these recycled components can be used to prepare DONs over multiple rounds with properties that are nearly identical to those prepared with pristine components. We recycle scaffold strands over multiple rounds by reprogramming them into completely new structures using a new set of staples to compete out staples from previously folded structures. Additionally, we recycle staple strands by recovering excess strands from DON folding and using them in subsequent folding reactions over multiple rounds, thus significantly reducing waste. We used Transmission Electron Microscopy (TEM) and fluorescence measurements to show recycled scaffold and staple strands can produce structures that are visually indistinguishable from pristine DONs and exhibit the same mechanical properties. Our results establish approaches for sustainable manufacturing of DONs, further paving the way for clinical and commercial applications through cost-effective and environmentally considerate methods. (37,38)

## Results and Discussion

### Scaffold Reprogramming

To recycle the scaffold component, our approach is to reprogram the scaffold, which is inspired by prior efforts studying competitive self-assembly of DONs where two sets of staple strands compete for the same scaffold. (34,39) In particular, one study showed that, in the presence of two different sets of staple strands, having a significant excess of one staple set leads to folding of just the one structure corresponding to the larger staple concentration. (34) We reasoned that, if those conditions are realized *in situ* by melting an already folded DON in the presence of an alternative set of staple strands, a structure could be refolded into a completely new device.

Towards this end, we developed a procedure to recycle scaffold from a folded DON by “reprogramming” it into a different DON. We achieved this by melting the structure and competing out incumbent staples with an excess of new target structure staples. This scaffold reprogramming strategy allows re-use of the same scaffold strands to create different structures over multiple folding cycles (Figure 1). To realize scaffold reprogramming, we combine DONs with target structure staples and apply a 95°C heating step before proceeding with typical folding temperature profiles (Methods, Table S5).

**Figure 1:**
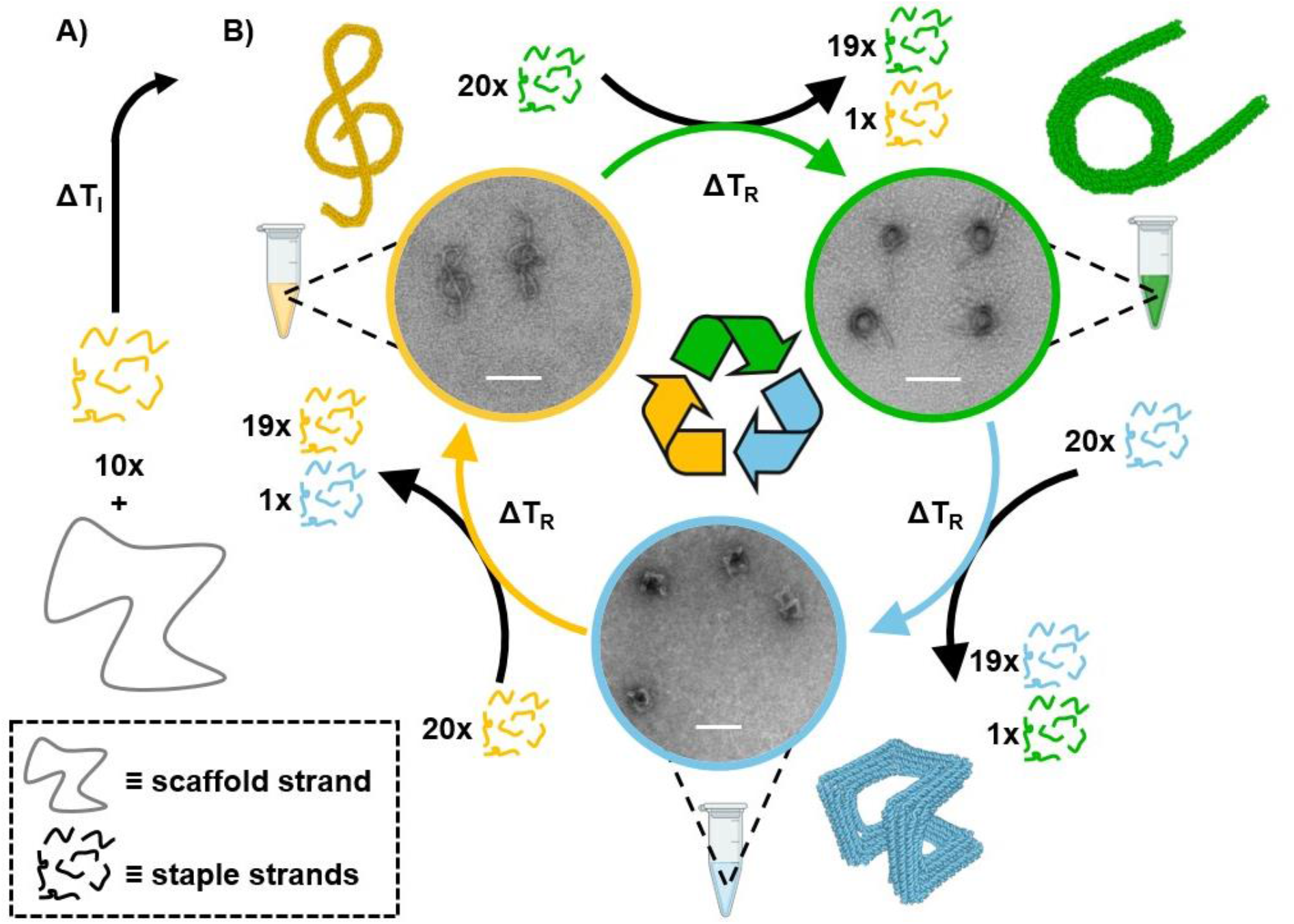
Schematic of scaffold recycling through reprogramming. A) Folding of a DNA origami structure through initial thermal annealing (ΔT_ι_). B) Reprogramming one structure into a new structure through competitive binding of target staple strands, using thermal annealing for reprogramming (ΔT_R_). TEM micrographs show representative structures after reprogramming. TEM images and oxDNA models are used to highlight the different shapes and confirm the experimental results agree with simulated structures. Scalebars: 100 nm

As a first demonstration of reprogramming a DON, we chose to use three recently reported designs (40): the G-Clef, NuSpring, and Hilbert structures (Figure 1). We selected these DONs to cover a broad spectrum of structurally distinct designs. We implemented the scaffold reprogramming approach (see Supplementary Information for details), starting with G-Clef structures folded from pristine scaffold and staples (Figure 1A). The pristine G-Clef structures were then reprogrammed into NuSpring structures; the reprogrammed NuSprings were then reprogrammed into Hilbert structures; and finally, the reprogrammed Hilbert structures were reprogrammed back into G-Clef structures, completing a full scaffold reprogramming cycle (Figure 1B). Successful scaffold reprogramming was verified by TEM (Figure 1B insets).

In addition to directly evaluating the shape of reprogrammed structures, we also compared them to pristinely folded structures for all three designs. Figure 2A and Supplementary Figures S1-3 show TEM images of the reprogrammed NuSpring, Hilbert, and G-Clef structures, again revealing well-folded structures similar to the pristine structures. Recycled and pristine structures also matched the shape of structures predicted by oxDNA simulation (Figure 2A left). Gel electrophoresis analysis showed that reprogrammed structures exhibit similar electrophoretic mobility (Figure S5). We observed a small shift in gel mobility between pristine structures and reprogrammed structures with reprogrammed structures running slightly slower. A similar difference in mobility has previously been observed between polyethylene glycol (PEG) purified samples (41) and unpurified samples and may be due to trace amounts of PEG remaining from the reprogramming process (Figure S5A, Figure S7).

**Figure 2:**
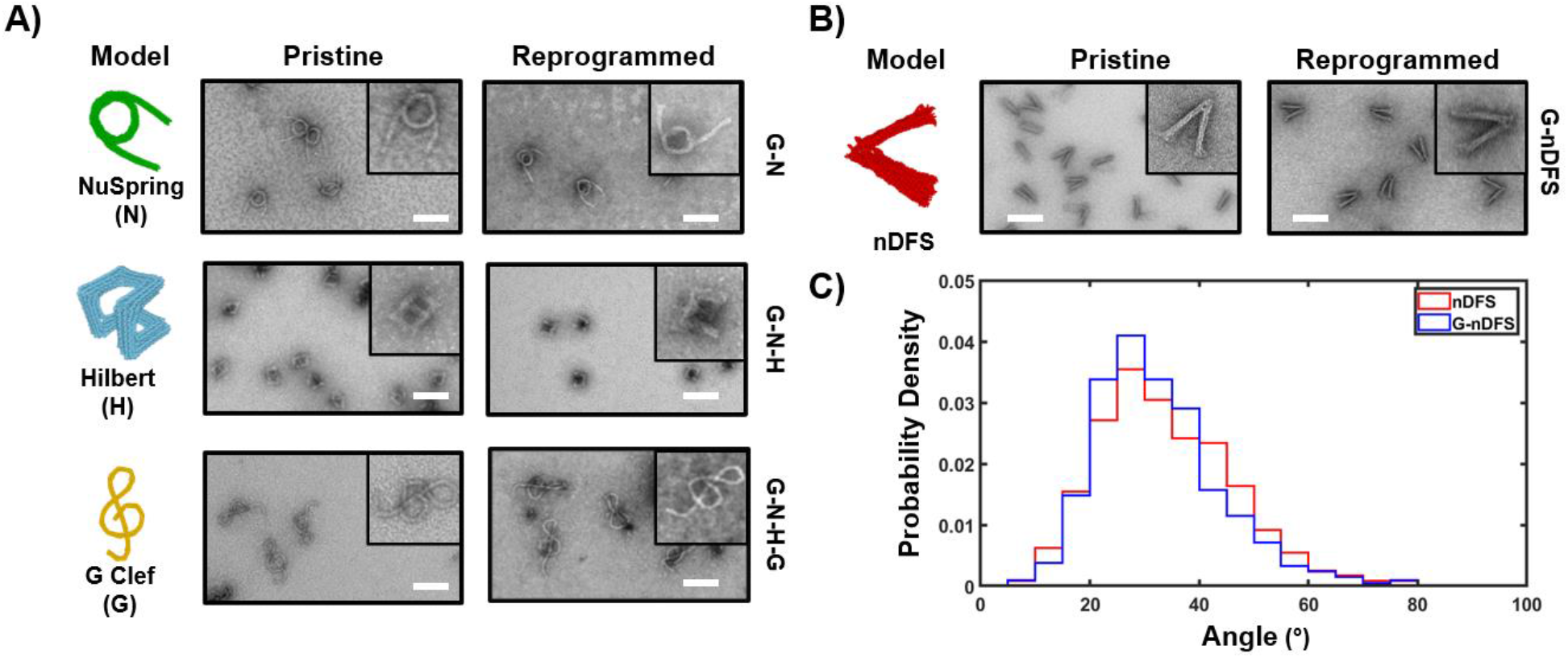
Analysis of scaffold reprogramming A) Imaging analysis of cyclic scaffold reprogramming of three structures and visual comparison with pristine examples through TEM imaging. B) - C) Quantitative analysis of scaffold reprogramming nDFS device through comparison of pristine and reprogrammed TEM micrographs (B) and measured angle conformation (C). Scalebars: 100 nm

We quantified the yield (i.e. fraction of well-folded structures) of the scaffold reprogramming process by gel densitometry (Figure S5A and Table S1). The first round of cyclic reprogramming (G -> N) showed 86% yield of well-folded structures compared to 88% for the pristine NuSpring sample. After the final reprogramming step (GNH -> GNHG), the recycled structures showed a yield of 81% compared to 93% yield for the pristine G-Clef.

While the yields remain relatively high, the drop in yield, especially in the GNH -> GNHG step, may be due to early rebinding of incumbent strands which have high melting temperatures. Analysis of the melting temperature of continuous dsDNA domains for the three different structures shows that the Hilbert structure has more domains with higher melting temperature (>60°C) and a higher maximum dsDNA domain melting temperature compared to the other two structures (Figure S6). During the thermal annealing process of reprogramming, it is possible that these incumbent staples with high melting temperature domains will rebind stably early in the thermal ramp after the initial 95°C melting step but before the target structure staples corresponding to the same domain can bind, thus making reprogramming less efficient.

To evaluate staple strand incorporation in the reprogramming process, we fluorescently labeled one G-Clef staple strand to track this staple when reprogramming to the NuSpring structure. (Figure S7). This staple was chosen to be the strand with the highest calculated melting temperature for a single binding domain. As such, we would expect this to be one of the most challenging incumbent strands to compete out during reprogramming. Fluorescent imaging of the reprogrammed NuSpring structures showed no detectable signal from the labeled strand, indicating that the incumbent staples are effectively out-competed by the NuSpring staples during reprogramming (Figure S7).

While single-staple labeling provided us a way to “spot check” that staples are removed, we cannot easily confirm whether all the incumbent staples are effectively competed off from the scaffold. And while the reprogrammed NuSpring, Hilbert, and G-Clef structures all exhibit the intended new shape, we also wanted to test whether reprogrammed structures exhibit similar properties to pristine examples. To this end, we chose to quantify the mechanical properties of a DON previously used for molecular force measurements referred to as the nanoscale DNA force spectrometer, or Ndfs (42,43). The mechanical properties of the nDFS, measured in terms of the angular distribution of the hinge-like design, are highly sensitive to staples at the vertex. We previously showed that substitution of a few staples leads to large shifts in angular distributions. (42)

We started by folding a pristine version of the G-Clef and then reprogrammed it into a variant of the nDFS referred to as nDFS-C35 (Figure S8A) (42). We characterized these reprogrammed structures by TEM imaging and agarose gel electrophoresis and observed successful refolding into structures indistinguishable from pristine samples (Figure 2B, S4, and S5B). The angular distributions of the two structures closely agreed, indicating reprogrammed devices can maintain functional properties in addition to the target structure (Figure 2C).

### Staple Recovery

While this reprogramming approach enables reuse of scaffold, a major source of waste from DON fabrication is the large excess of staple strands. DONs are usually folded with 10-fold excess of staples leading to 90% of staples being wasted. To improve material efficiency of DON folding, we developed a protocol to recover excess staples for re-use in subsequent folding reactions. Our approach, depicted schematically in Figure 3A, builds off of the purification approach of centrifugal precipitation in the presence of PEG, which is widely used to remove excess staple strands in DNA origami fabrication. (41) This process yields a purified structure pellet, which can be resuspended into a desired buffer, and a supernatant containing PEG and the remaining excess staples. This supernatant is typically discarded as waste. To re-use these excess staples in subsequent folding reactions, we first tested using the supernatant directly (i.e. in PEG solution) in a folding reaction, adjusting the staple concentration to the typical 10-fold excess relative to scaffold. This approach led to pervasive aggregation as observed by gel electrophoresis, which precluded structural analysis and effective folding of new DONs (Figure S9-11).

**Figure 3:**
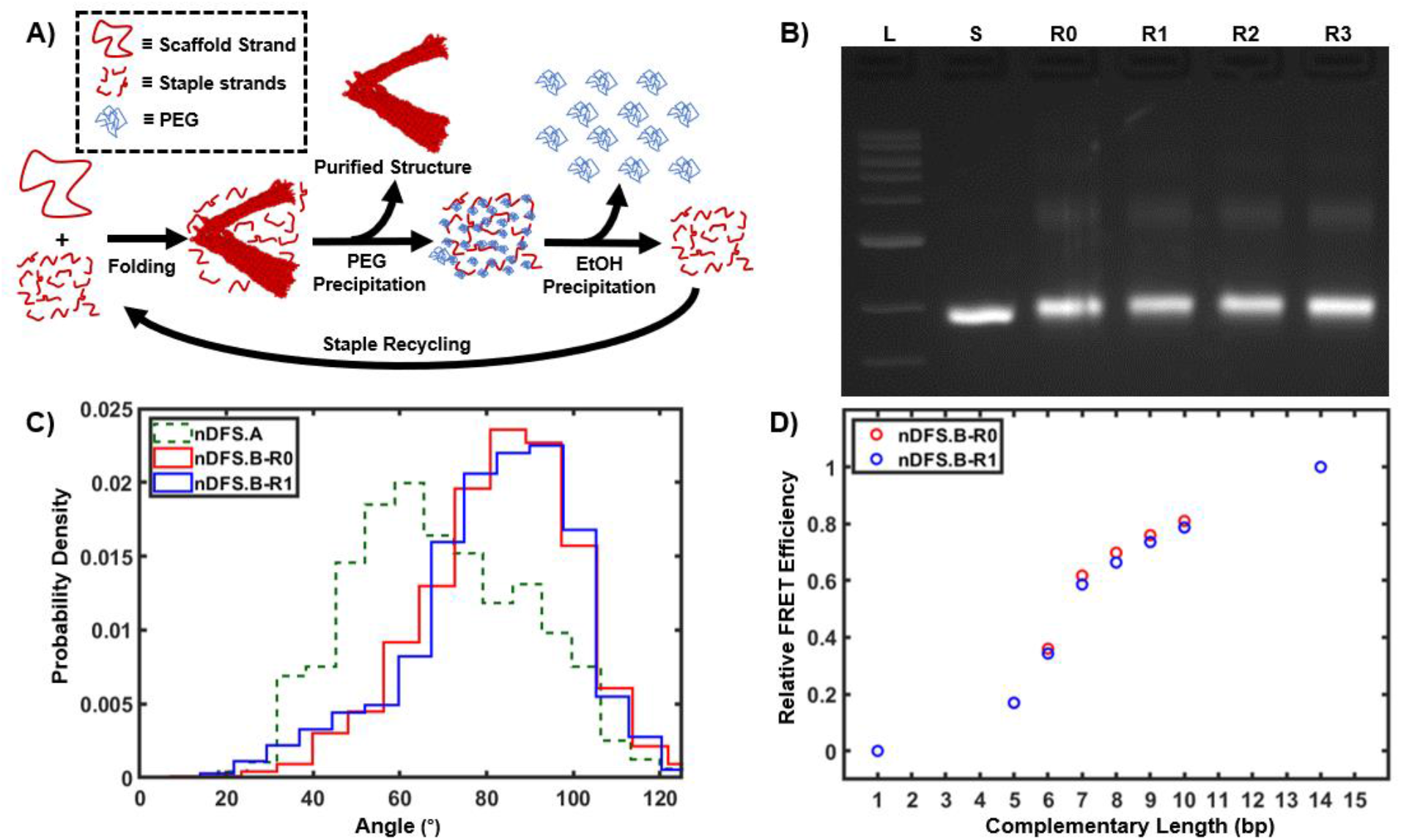
Recycling through staple recovery. A) Schematic representation of the staple recovery process through ethanol precipitation of supernatant from PEG precipitation. B) Agarose gel electrophoresis of nDFS.B structures folded using recycled staples over three rounds. C) Comparison of TEM-measured angle distributions of pristine (R0) and once recycled staples (R1) nDFS.B structures with nDFS.A distribution shown for comparison. nDFS.A data was previously reported in (42) D) Relative FRET Efficiency of nDFS.B structure with labeled latch system for both nDFS.B-R0 and nDFS.B-R1 structures. Measurement uncertainty was smaller than the datapoints and has been reported in Table S3.

To separate the staples from the PEG solution, we used ethanol to precipitate the staple oligonucleotides. We found that staples could be precipitated efficiently by incubating the PEG supernatant with a refrigerated mixture of 90% ethanol and MgCl_2_ (final concentration of 11 mM). This ethanol solution was mixed at a ratio of 2:1 with the PEG supernatant and centrifuged to yield a pellet of DNA staples. The ethanol and PEG supernatant was removed, and the precipitated staples were then resuspended in 1x TE (10 mM Tris,1 mM EDTA) for use in future folding reactions. Again, we selected the nDFS hinge to demonstrate the efficacy of our staple recovery protocol, this time choosing a different vertex design that relies on staple strand incorporation (nDFS.B) (Figure S8A). (42) This version exhibits a higher average hinge angle due to incorporation of a few staple strands at the hinge vertex compared to the nDFS.A, where the scaffold connections at the vertex remain single-stranded. Hence, in addition to TEM visualization of correct structure folding using recovered staples, the shift of the angular distribution relative to nDFS.A is an additional indication of the incorporation and fidelity of recycled staples near the vertex.

We compared pristine nDFS.B structures (R0) and nDFS.B structures folded using recovered staples (RN where N = cumulative number of staple recovery rounds) by multiple methods. We demonstrated the ability to recover and reuse staples over three rounds of folding. Samples R0 through R3 all exhibited similar mobility in agarose gel electrophoresis experiments (Figure 3B). Additionally, we used TEM and ensemble fluorescence methods to compare functional properties of the nDFS.B structures, pristinely folded or folded with recovered staples. We compared the angular distributions of R0 and R1 structures by TEM (Figure 3C, Figure S12). The nDFS.A angular distribution is also shown in a dashed outline for reference. The nDFS.B angle distribution could revert to the nDFS.A distribution if either the vertex staples were not recovered during the recycling process or there were defects in staples that led to poor incorporation. The angular distributions showed a clear shift relative to nDFS.A and good agreement between the pristine and recycled structures, suggesting structures folded with recovered staples exhibit similar mechanical properties (Figure S12). As with scaffold reprogramming, we quantified the yield of folding through gel analysis and found that folded structure yield was similar to that of pristine structures maintaining a yield of approximately 90% through three rounds of recycling (Table S2).

To more directly quantify the intended function of the nDFS devices, which is to apply forces to molecular interactions, we repeated a previously reported set of experiments designed to test the stability of a base-pairing interaction placed between the two arms. (43) We added overhanging staples protruding from the inner faces of the top and bottom arms so that they could bind via base-pairing to latch the two arms together. We used a fixed length for the top arm overhang and a bottom staple of variable length to tune the length of hybridization, thus allowing us to control the stability of the bound state. Binding of the overhangs also requires the hinge to be latched to a smaller angle, and we previously showed that the hinge applies forces on the scale of ∼10 pN, shifting the interaction to the dissociated state. (43) We incorporated fluorophores as a readout of latching (Figure S8B) such that when the overhanging strands are hybridized, the fluorescent labels undergo Forster Resonance Energy transfer (FRET). This allowed us to measure the average closing of the nDFS.B hinge by measuring FRET efficiency, (44) while varying the strength of the latching interaction. Figure 3D compares the average FRET measurements for staple-recycled (R1) and pristine (R0) nDFS devices across different complementary overhang lengths. To correct for cumulative photobleaching of recycled fluorescent staples, each dataset was min-max normalized using the mean 0 and 14 bp nDFS.B measurements. The staple-recycled and pristine samples show excellent agreement across the range of complementary lengths tested, demonstrating that staple-recycled samples apply similar tensile forces to overhanging strands and thus have comparable functionality for the intended application of force measurements.

In addition to this mechanical function, we also tested the ensemble behavior of a DON in a biologically relevant setting through performing small molecule drug (Daunorubicin) loading and release experiments with a DON that has been studied extensively before. (19,45) The drug binds to the DON through intercalation, and hence the loading and release behavior should be sensitive to the correct incorporation of staple strands. (46) We observed similar loading levels and release profiles of Daunorubicin, an anti-cancer drug, for DON folded with pristine and recovered staple strands (Figure S13).

### Minimal Waste Fabrication

The scaffold reprogramming and staple recycling procedures described above can be combined to reuse both the scaffold and staple strands of a folded structure allowing for minimal waste fabrication of origami structures. When following our protocol for scaffold reprogramming, the PEG precipitation supernatant can be retained, and the remaining 19x excess of target device staples can be purified through ethanol precipitation and used for subsequent folding or reprogramming reactions. To demonstrate this minimal waste fabrication, we started by folding NuSpring and G-Clef structures using pristine staples and scaffold. These structures were then purified through PEG precipitation, and then the excess staples were recovered through EtOH precipitation of the PEG supernatant. These recovered staples were then combined with the folded structures of the other type (e.g. NuSpring recovered staples with G-Clef purified structures and vice versa) and reprogrammed using the same thermal protocol described above, as schematically depicted in Figure 4A. This allows for the synthesis of new DONs without using any pristine staple or scaffold materials. These minimal waste structures were verified by agarose gel electrophoresis and TEM imaging (Figure 4B, C and S14-15), which both indicated well-folded recycled structures.

**Figure 4:**
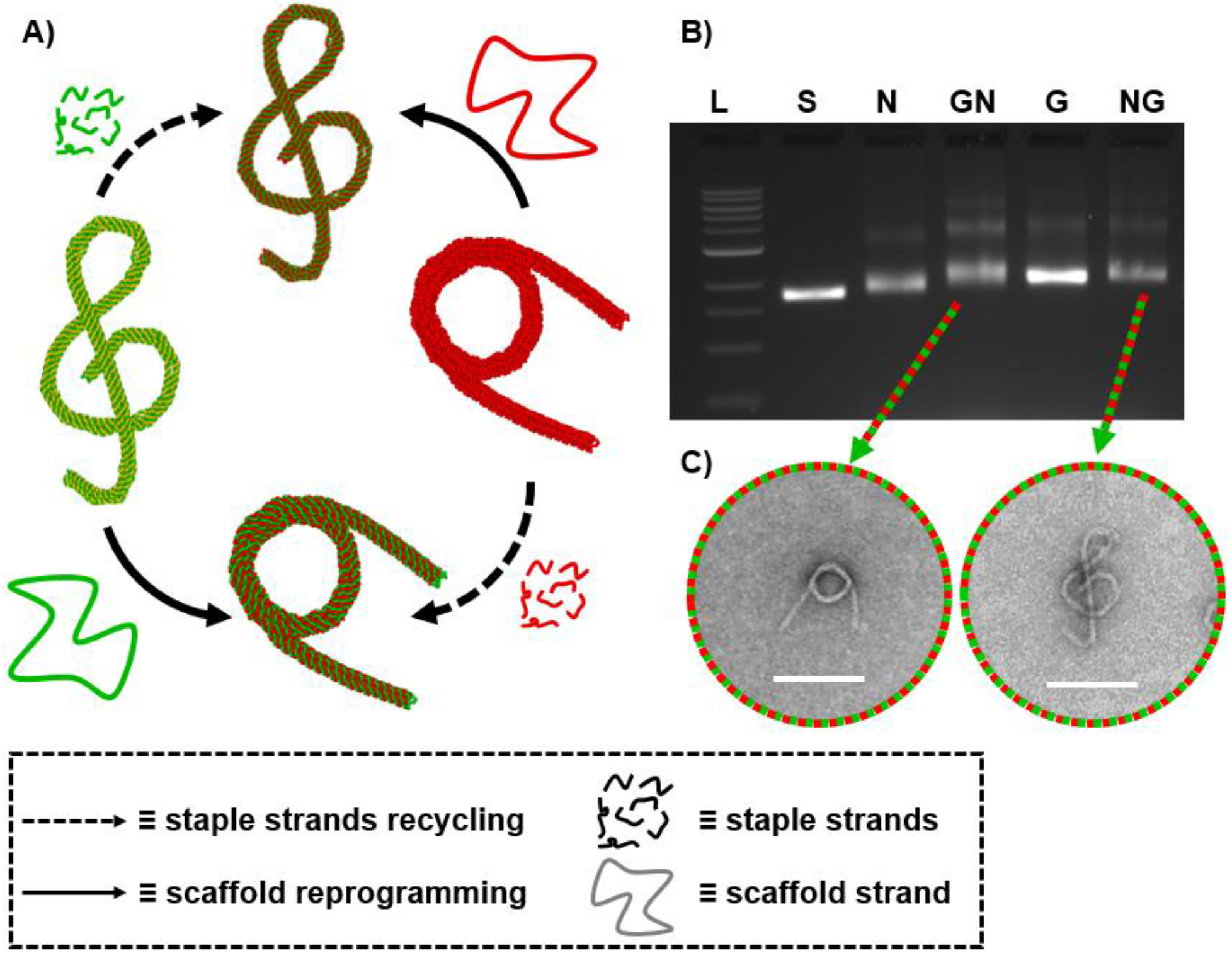
Minimal waste fabrication. A) Reprogramming folded G-Clef and NuSpring structures into each other, using excess staple strands recovered after folding from the respective structures. B) Agarose gel electrophoretic analysis of pristine and reprogrammed devices using the minimal waste fabrication protocol (L = 1kb ladder, S = p8064 scaffold, N = pristine NuSpring, GN = G-Clef reprogrammed into NuSpring, G = pristine G-Clef, NG = reprogrammed G-Clef). C) representative TEM images of the reprogrammed devices. Scalebars: 100 nm

## Conclusions

We introduced significant advances toward sustainable DNA origami manufacturing by recycling the two constituent DNA materials: scaffold and staple strands. We demonstrated reprogramming of scaffold DNA from a folded structure into a completely new design, and recovery of excess staple strands for use in subsequent folding reactions. We determined the yield of recycled DONs to be between 81% and 91% and verified both recycling approaches retain functional properties of dynamic devices. We further combined these methods to demonstrate effective folding of DONs from completely recycled DNA materials, establishing a route for maximizing material-use efficiency and achieving an extended life cycle in materials used for structural DNA nanotechnology. (37,38,47) Use of these recycling strategies may likely require some case-specific validation of structure and properties comparable to established recycling protocols. Further optimization of reprogramming could yet improve recycling yields, for example, purposeful design of staples to facilitate recycling by avoiding domains with high melting temperatures. (35) Additionally, these protocols could be expanded to allow recycling of, and into, multi-scaffold devices, to meet current advances. (13,48)

While most DNA origami research has been conducted in an academic setting, there has been a recent increase in the number of DNA origami-related patents filed, indicating forthcoming industrial applications. (49) As research efforts are scaled up towards commercial applications, recycling of origami through scaffold reprogramming and staple recovery should be considered to extend the life of expensive DNA reagents and minimize the waste generated through fabricating DONs. Staple recovery may prove especially useful for DONs that use expensive or difficult to prepare nucleic acid molecules such as staple strands with costly modifications e.g. fluorophores, groups for chemical conjugation, or synthetic RNA molecules. (50) The recycling techniques presented here have the potential to facilitate future applications that require especially large amounts of DONs such as assembly of 3D materials where the bulk properties are of interest or applications in cell, tissue, or biological systems, where challenges remain including biocompatibility, stability, and scaling up fabrication and purification processes. (19,51–53). Furthermore, these methods may be impactful to laboratories with limited resources or budgets to maximize the use of materials for DNA origami self-assembly. Even as commercial DNA synthesis is further optimized, it will be important to develop and implement efficient, cost-effective use of nucleic acid reagents and to reduce the waste associated with their use. (15,54,55) Moving forward, this work can be integrated synergistically with other efforts to reduce costs of scaffold and staple production ultimately advancing DNA nanostructure manufacturing. (18,56,57)

## Supporting information

Recycling Sequences

## Supporting Information

Methods, Uncropped TEM images of recycled DONs, gel electrophoresis of recycled DONs with densitometry analysis thereof, and additional experimental details. (PDF) DNA origami staple sequences. (Excel)

## Author Contributions

This project was conceived by W.G.P. Experiments were led by M.J.N. and Y.W., including optimization of staple strand recovery and reprogramming of DNA origami structures. Y.W. wrote the MATLAB code to analyze melting temperatures. W.G.P. performed early proof-of-principle experiments. N.V. performed auxiliary experiments.

M.G.P. and C.E.C. acquired funding; M.G.P., C.E.C. and W.G.P. supervised the study and helped with data interpretation. M.J.N. and W.G.P. prepared figures; M.J.N. and W.G.P. wrote the manuscript with input and feedback from C.E.C. and M.G.P. All authors reviewed and approved the final version. M.J.N. and Y.W. contributed equally.

## Notes

The authors declare no competing financial interest.

## Acknowledgments

We would like to thank all members of the Castro and Poirier groups for helpful discussions, feedback, and the positive and fruitful atmosphere. We acknowledge resources from the Campus Microscopy and Imaging Facility (CMIF), The Ohio State University.

