## Supplementary Information for "Recycling Materials for Sustainable DNA Origami Manufacturing"

### Supporting Information

### Methods

#### DNA Origami Design and Folding

DONs were designed using caDNAno and magicDNA. (1–3) Design files are available at www.nanobase.org (4). DONs were folded by combining 10 nM of 8064 nt scaffold isolated from M13mp18 bacteriophage with 100 nM of each staple strand of the corresponding structure in a solution containing 1 mM EDTA, 5 mM Tris, 5 mM NaCl, and 6 mM $MgCl_2$ (with the exception of nDFS samples which used 8 mM $MgCl_2$). This solution was divided into 50 µL aliquots and then subjected to a thermal ramp (described in Table S4) in a BioRad C1000 thermocycler.

#### Transmission Electron Microscopy (TEM)

DNA origami structures were imaged by TEM following previously established protocols (3). Purified DNA origami structures were applied onto glow-discharged Formvar-coated copper EM grids (Ted-Pella). Samples were diluted into 1 nM and incubated for 4 minutes before the excess sample was removed with filter paper. Next, samples were stained with 1% Uranyl acetate (SPI Supplies) or 2% Uranyl formate (Electron Microscopy Sciences). Imaging was performed with an FEI Tecnai G2 Spirit, using an acceleration voltage of 80 kV at the OSU Campus Microscopy and Imaging Facility.

#### Atomic Force Microscopy

AFM imaging was performed as previously described. (5) Briefly, samples were deposited onto a freshly cleaved mica surface (Plano GmbH) and adsorbed for 2 minutes at room temperature. Subsequently, the mica was carefully rinsed with ddH2O and then dried using compressed air. Scanning was performed using the ScanAsyst in Air mode, with a Bruker BioScope Resolve AFM, equipped with a Nanoscope V controller. Silicon Nitride probes with a nominal spring constant of 0.4 N/m and sharpened pyramidal tips (Bruker) were used. Typically, scan rates around 1 Hz were used with scan sizes up to 3 µm. For larger scan sizes, the scan rate was lowered to reduce imaging artifacts. Images were analyzed using Nanoscope Analysis v1.9.

#### Ensemble FRET

Measurements were performed at 1 nM DON concentration in a solution containing 11 mM $MgCl_2$, 45 mM Tris, 45 mM boric acid, and 1 mM EDTA. Experiments were done using a Horiba Instruments FluoroMax4 spectrofluorometer using a 45 µL quartz cuvette. FRET efficiency was calculated by the RatioA method using the emission spectra of Cy3 and Cy5 fluorophores. (6)

#### oxDNA Simulation

We performed oxDNA simulations, allowing us to compare computational and experimental results. (7) Briefly, caDNAno design files were converted into oxDNA input files using tacoxDNA. (8) Simulations for the nDFS were first relaxed for 1E+7 steps, and subsequently run for 1E+8 steps, using default parameters from oxDNA.org. Averages over the full trajectory were used to prepare graphics in oxVIEW. (9,10)

**PEG Purification**

Purification of DNA origami nanostructures was performed by following a published protocol. (11) In Polyethylene glycol (PEG) purification, folded samples were mixed with an equal volume of solution containing 15% PEG (Molecular Weight 8000), 500 mM NaCl, 5 mM Tris. Then the mixture was centrifuged at 16,000 g for 30 min. After centrifugation, the folded structures were precipitated into a pellet at which point the supernatant, which contains PEG and excess staples, were removed by pipetting (and saved for staple recovery when applicable). Then the remaining pellet containing purified structures was resuspended into a suitable buffer by pipetting 50 times with the full buffer volume.

**Scaffold Reprogramming**

To recycle the scaffold strand through reprogramming, the starting structures were purified after folding following the above PEG purification protocol. Since these purified structures were combined with other ingredients (and thus diluted) for the subsequent refolding, a resuspension volume following PEG purification was chosen to ensure a structure concentration of roughly 100 nM to allow us to keep a consistent final reprogramming product concentration of 10 nM. All samples to be used for reprogramming were resuspended in a solution containing 10 mM Tris, 10 mM $MgCl_2$. With these purified structures in place of scaffold, folding reactions were prepared in a solution containing 10 nM of the purified starting structure (used in place of scaffold), 200 nM of each target staple strand, 1 mM EDTA, 5 mM Tris, 5 mM NaCl, and 6 mM $MgCl_2$. In the case of nDFS samples 18 mM $MgCl_2$ was used for reprogramming. Then folding reactions were divided into 50 µL aliquots and subjected to the reprogramming thermal annealing protocol described in Table S5.

**Staple Recovery**

Staples were recovered from the PEG supernatant described above in the PEG purification protocol. We then took this PEG supernatant containing the target structure staples and combined it with twice the supernatant volume of 90% ethanol stored at 4C and added highly concentrated $MgCl_2$ to bring the final $MgCl_2$ concentration to 10 mM $MgCl_2$. After mixing the ethanol and PEG supernatant by pipetting, the samples were spun down in a refrigerated (0°C) centrifuge at 16,000 g for 30 minutes to precipitate the staple strands. After centrifuging, the supernatant (containing ethanol and PEG) was removed by pipetting. The pellet was then resuspended in a buffer containing 10 mM Tris and 1 mM EDTA for use in future folding reactions, taking care to resuspend in a small enough volume to allow for a final folding reaction staple concentration of 200 nM.

**Agarose Gel Electrophoresis**

All gels were run in 1.5% agarose at 90 V for 90 minutes and pre-stained with Ethidium bromide. Gels were prepared using TBE buffer (45 mM Tris, 45 mM Boric acid, 1 mM EDTA) and 11 mM $MgCl_2$. Gel rigs were submerged in an ice water bath.

### Drug Loading and Release

Horse DNA nanostructure was folded according to the procedure in Halley 2016 (12) starting with the 7249 M13mp18 scaffold. The scaffold was placed in solution at 100 nM with a buffer containing 5 mM Tris, 5 mM NaCl, 1 mM EDTA, 18 mM $MgCl_2$, and oligonucleotide staple strands at a 10:1 ratio of staples to scaffold. The mixture was placed in a water bath at 70°C for 30 minutes to melt the DNA followed by another water bath at 52°C for 2.5 hours. The mixture was then placed in a fridge at 4°C overnight. The unpurified Horse DNA nanostructure was placed in equal volume amounts of 15% PEG 8000 (Sigma Aldrich, St. Louis, MO). After one round of PEG purification, origami precipitated into a pellet, while the supernatant was collected. The supernatant, containing excess staple strands, was subject to ethanol precipitation which would be used for the next Horse folding reaction. Horse nanostructures were suspended in PBS, pH 7.4, 2.5 mM $MgCl_2$. Horse DNA nanostructures were loaded with daunorubicin HCl (Santa Cruz Biotechnology, Dallas, TX) to yield solutions of 40 nM Horse and 50 μM daunorubicin. Daunorubicin absorbance was measured on a plate reader (SpectraMax M2, Molecular Devices, Sunnyvale, CA) at 480 nm. Daunorubicin-loaded Horse mixture and daunorubicin were added to Slide-A-Lyzer Mini Dialysis Devices (Thermo Fisher Scientific, Waltham, MA) and placed in a water bath of PBS (Gibco, Life Technologies) at 37°C with light stirring through magnetic stir bar. Daunorubicin release was observed by measuring the absorbance of the daunorubicin-loaded structure mixture on a NanoDrop One (Thermo Fisher Scientific, Waltham, MA) at 480 nm following the company's instruction manual.

### Melting Temperature Analysis

Temperatures were calculated using a custom MATLAB code to identify continuous domains in a DNA origami design and to calculate a melting temperature for that domain. The melting temperature was calculated using Tm=dH/(dS+R*log(Cts/4)) where Tm is the Melting temperature in Kelvin, Cts is the concentration of the two binding strands (taken to be 20 nM), R is the ideal gas constant, and dH and dS are the total entropy and enthalpy values for a given set of nearest neighbor base-stacking interactions in the strand taken from the "Unified" values in SantaLucia and Hicks (13).

**Supplementary Figures**

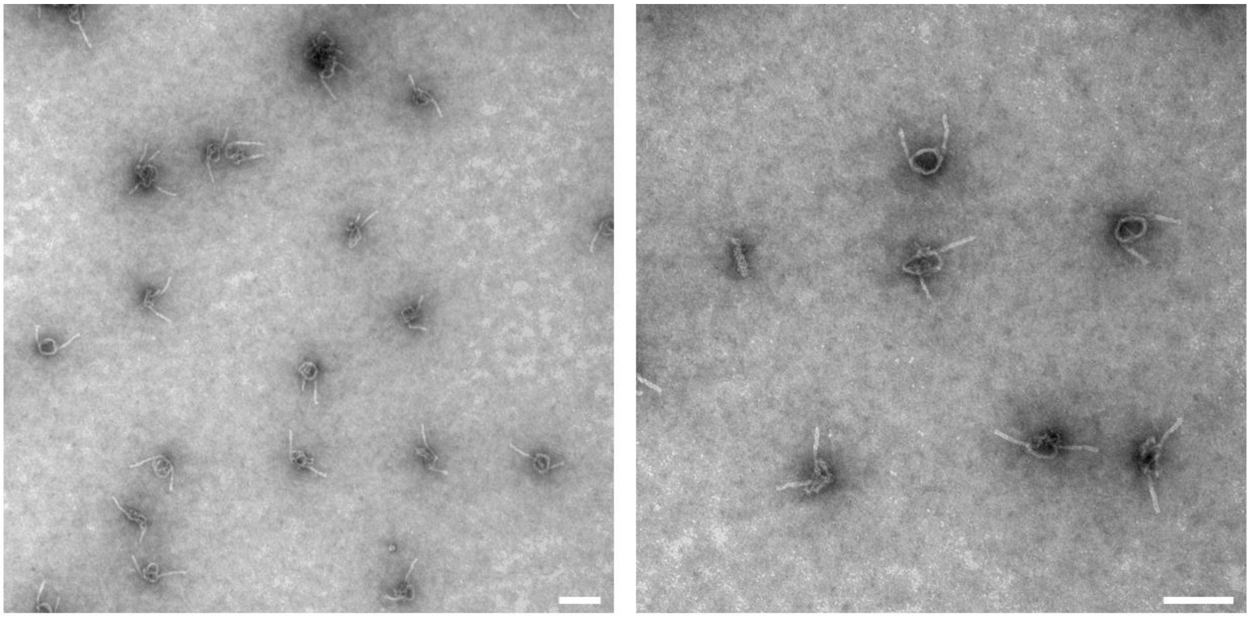

**Figure S1**: Uncropped TEM images of GN structures (G-Clef reprogrammed to NuSpring). Scalebars: 100 nm.

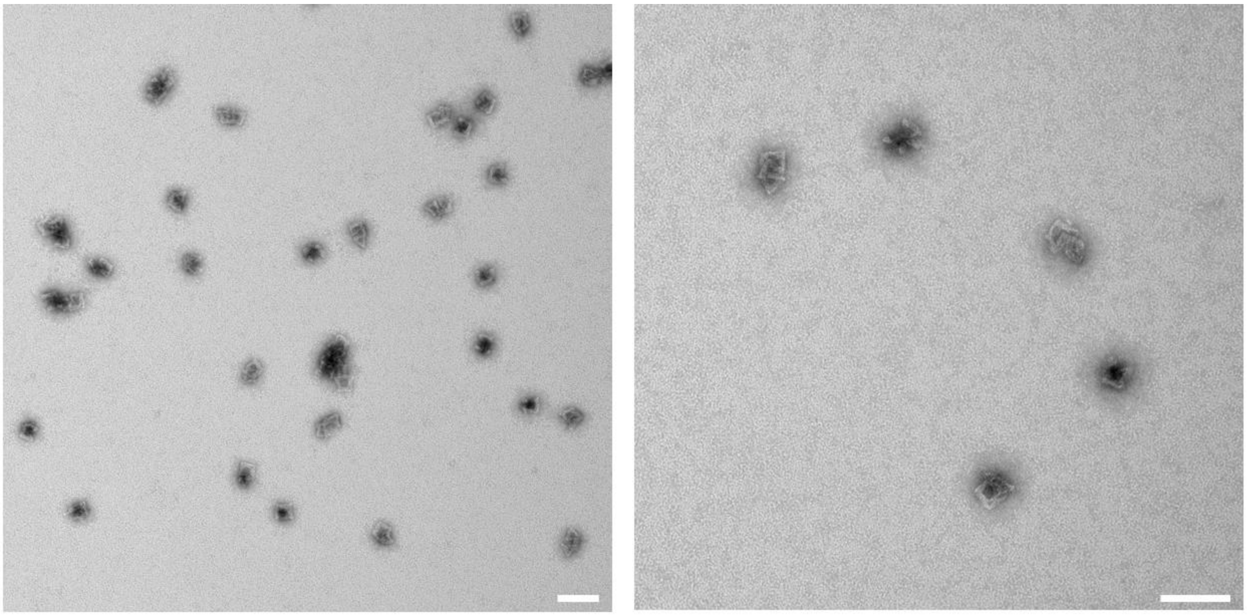

**Figure S2**: Uncropped TEM images of GNH structures (G-Clef reprogrammed to NuSpring reprogrammed to Hilbert). Scalebars: 100 nm.

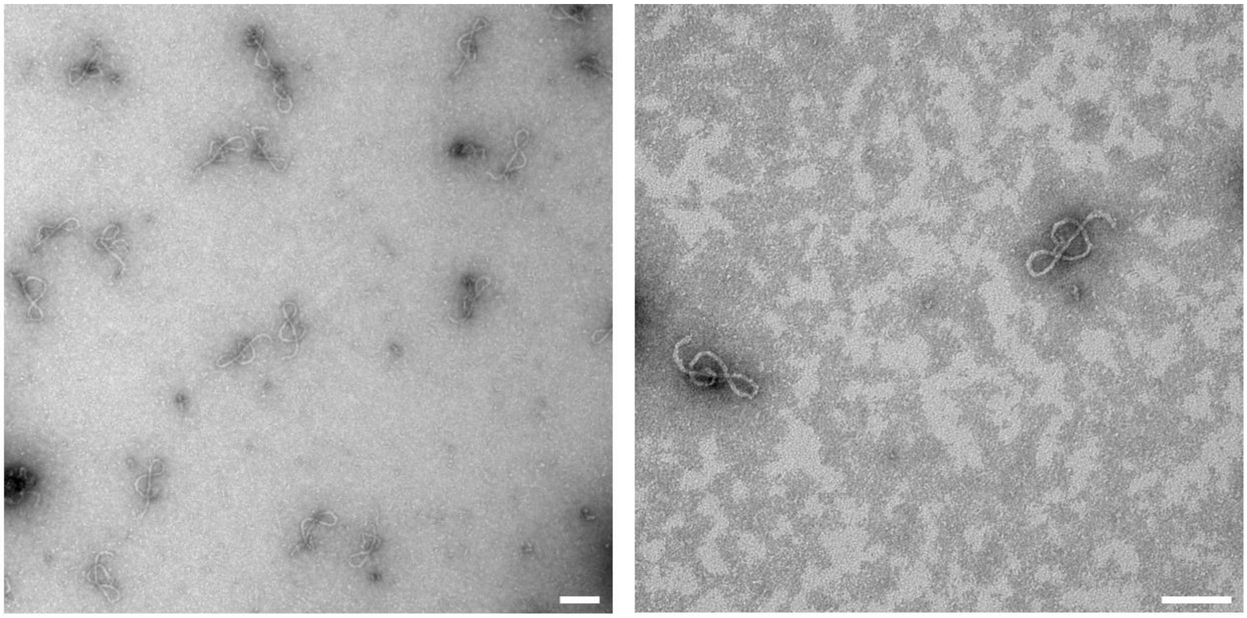

**Figure S3**: Uncropped TEM images of GNHG structures (G-Clef reprogrammed to NuSpring reprogrammed to Hilbert reprogrammed to Hilbert). Scalebars: 100 nm.

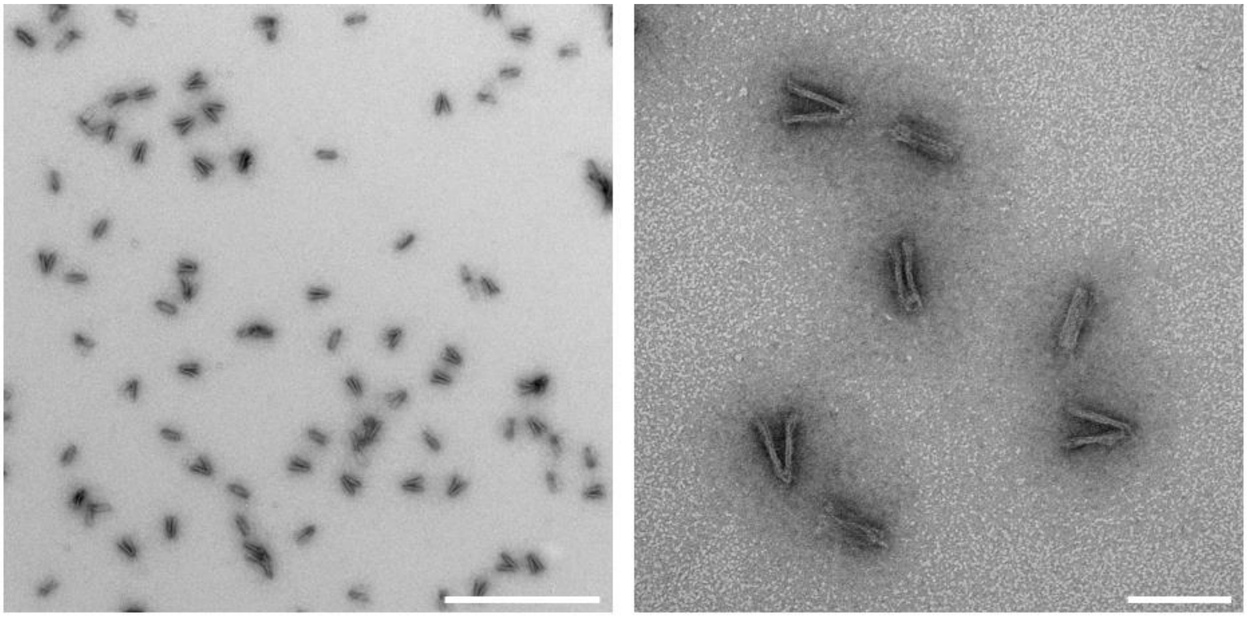

**Figure S4**: Uncropped TEM images of nDFS structures reprogrammed from G-Clef (G-nDFS). Scale bars represent 500 nm (left) and 100 nm (right) respectively.

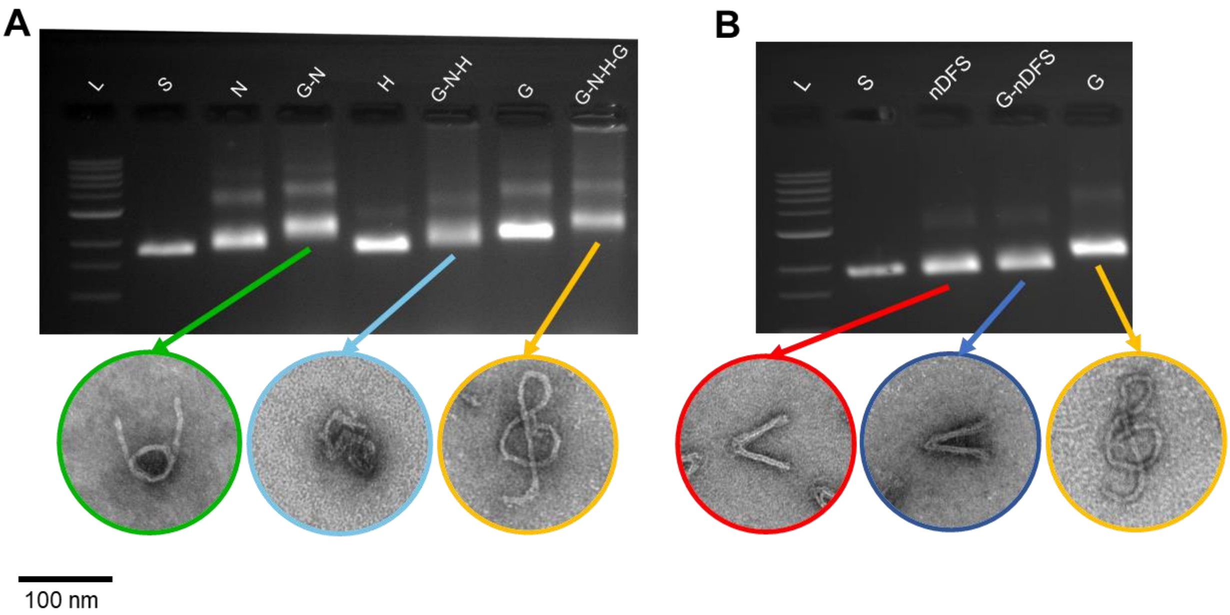


**Figure S5** (A) Agarose gel electrophoresis of cyclical scaffold reprogramming of NuSpring, Hilbert, and G Clef structures. These correspond to the same reprogrammed structure shown in Figure 2 A)**.** (B) Shows agarose gel electrophoresis of reprogrammed nDFS.C structures.

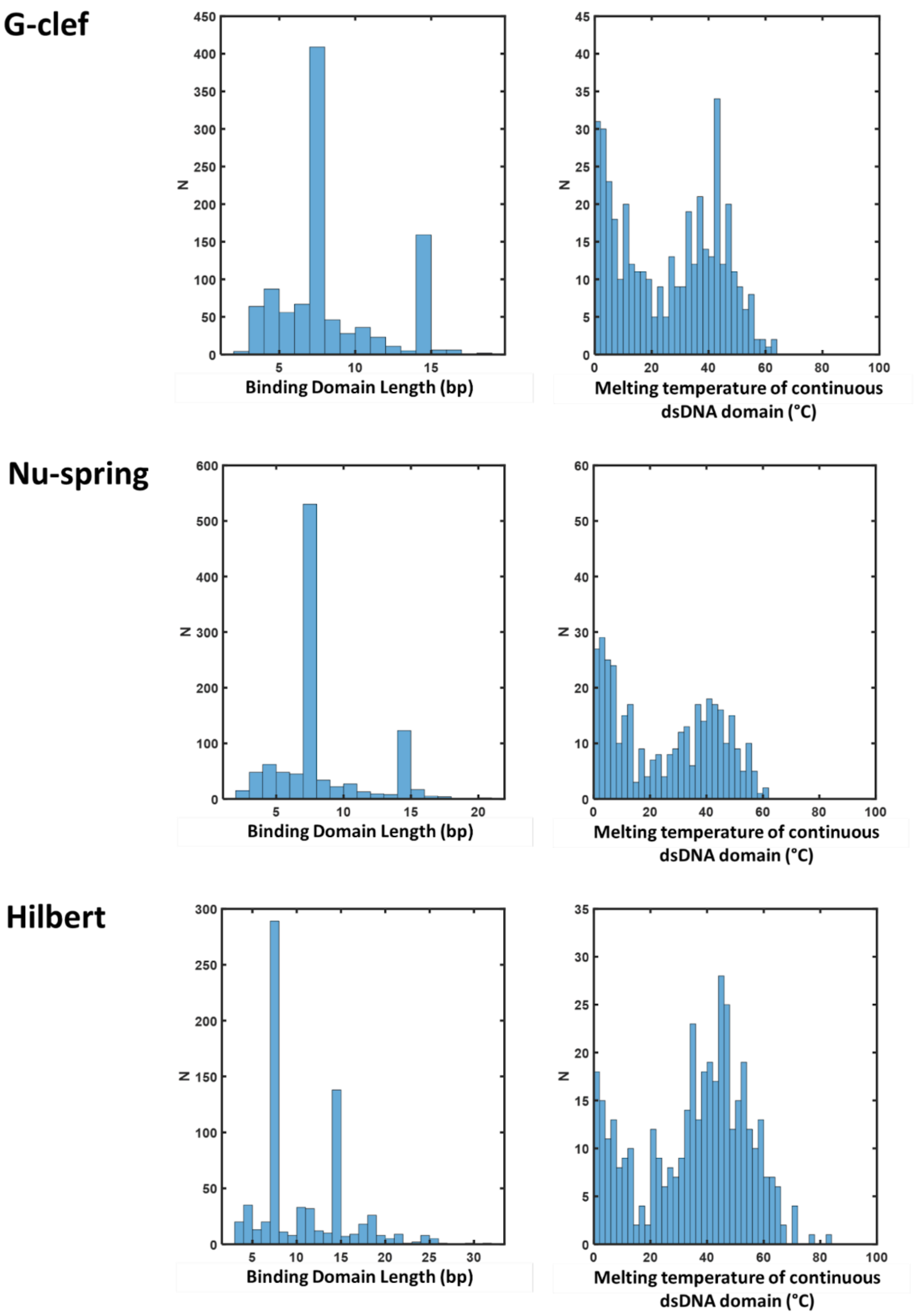


**Figure S6:** Distribution of binding domain lengths and melting temperatures calculated for every continuous (without crossovers or nicks) dsDNA domain for the three MagicDNA structures.

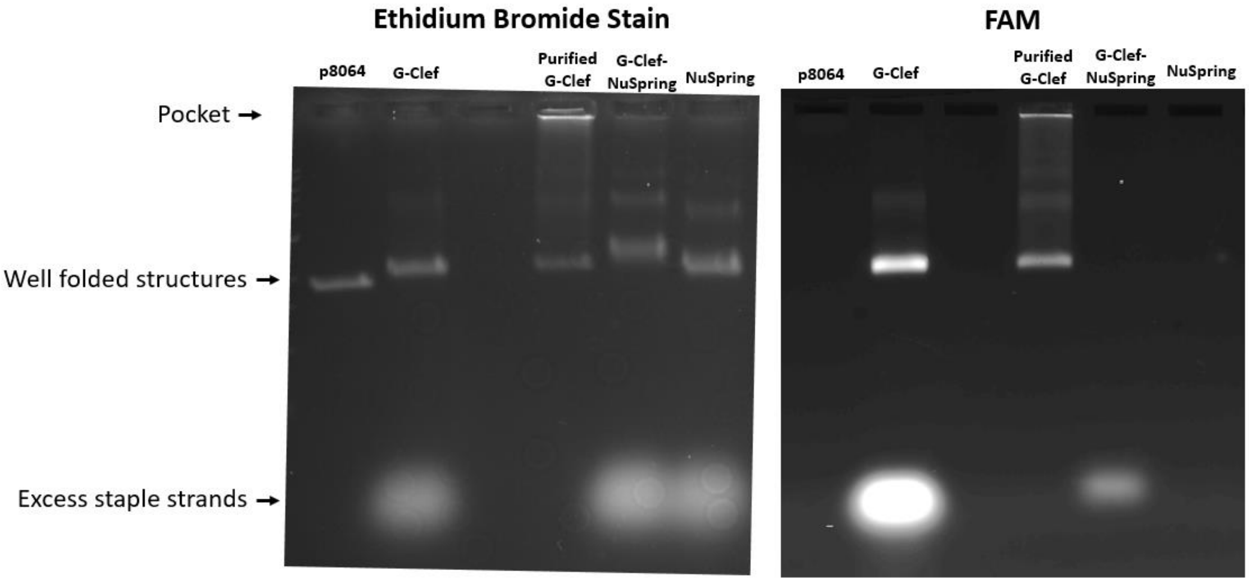


**Figure S7** Agarose gel electrophoretic analysis of G-Clef folded with a fluorescently (FAM) labeled staple (chosen for having the highest calculated melting temperature) and then reprogrammed into the NuSpring structure (G-Clef-NuSpring). The FAM channel shows no detectable signal in the “Well folded structures” band of the G-Clef-NuSpring suggesting incumbent staples are fully out-competed during the reprogramming process. The excess staple strand shows a small signal for G-Clef-NuSpring consistent with a 1x ratio of incumbent G-Clef staples.

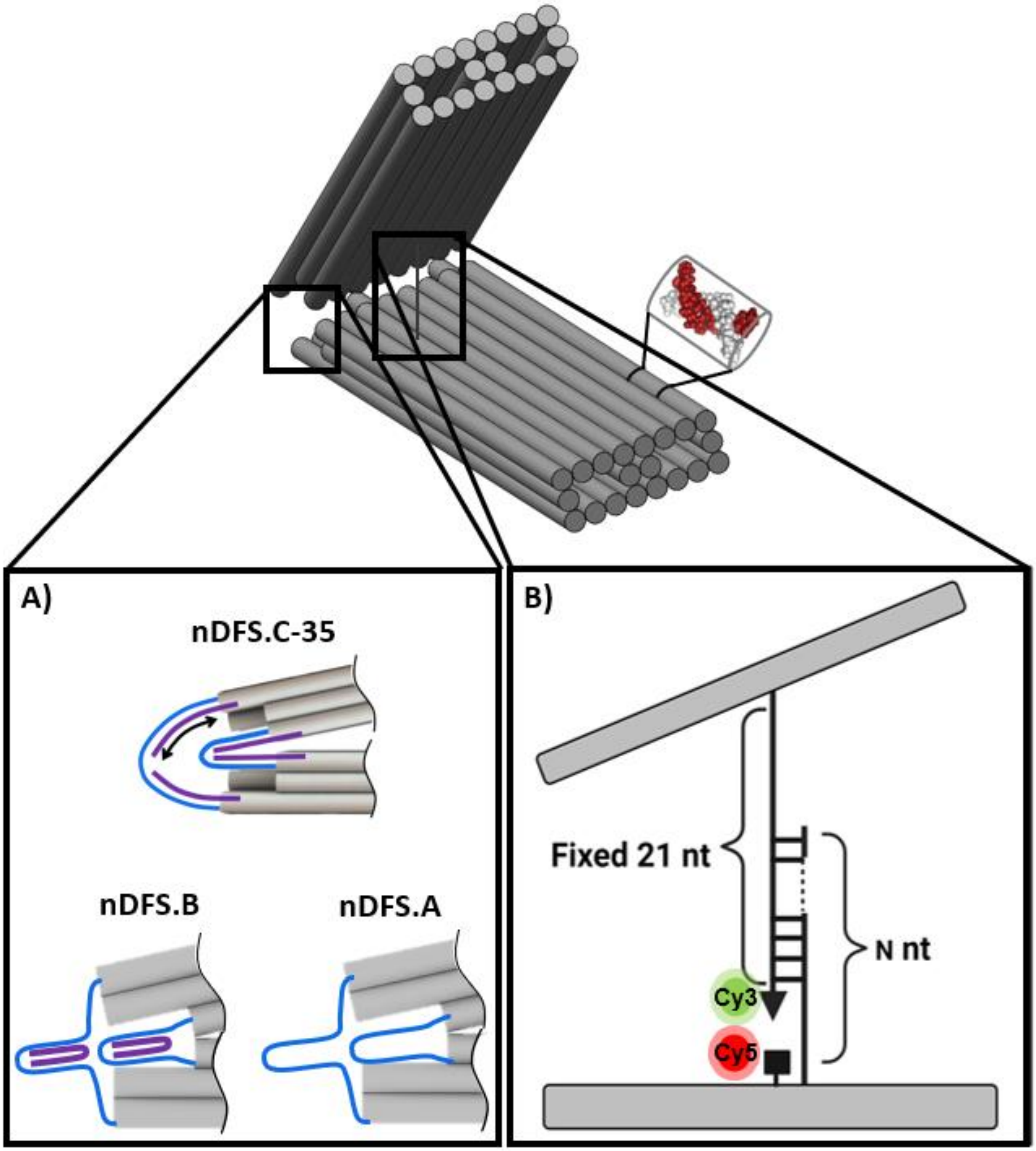


**Figure S8** Schematic of nDFS structure highlighting latching and vertex staples. A) shows the position of staple strands on the nDFS.C35, nDFS.B, and nDFS.A structures. The nDFS.C35 variant was used for all reprogramming experiments shown in Figure 2. The nDFS.B variant was used for staple recovery experiments and Figure 3 C shows the difference in angular distribution due to the presence of the staples strands which differentiate the nDFS.A and nDFS.B versions. B) shows the staple overhang and fluorophore labeling scheme used for the experiments in Figure 3 D).

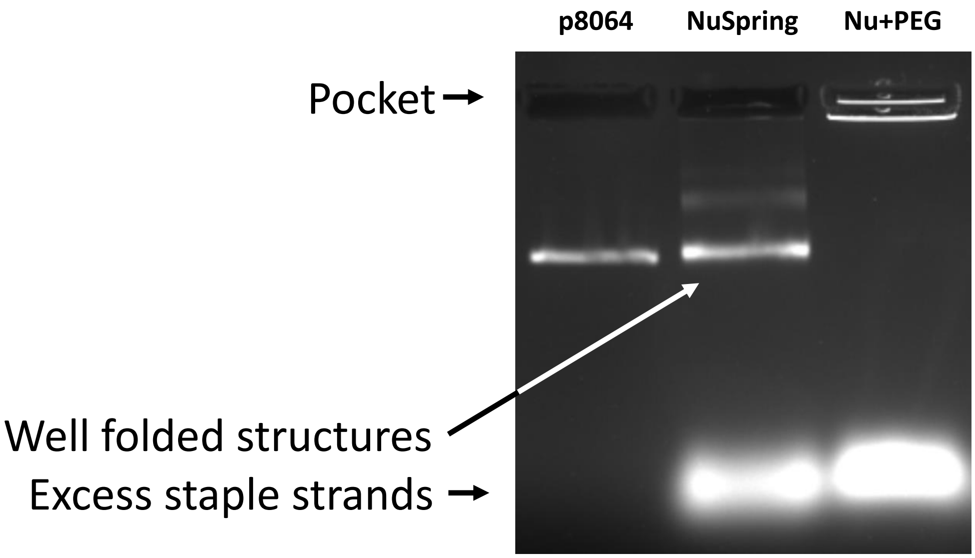


**Figure S9**: Agarose gel electrophoretic analysis of the NuSpring DNA Origami structure folded with or without PEG.

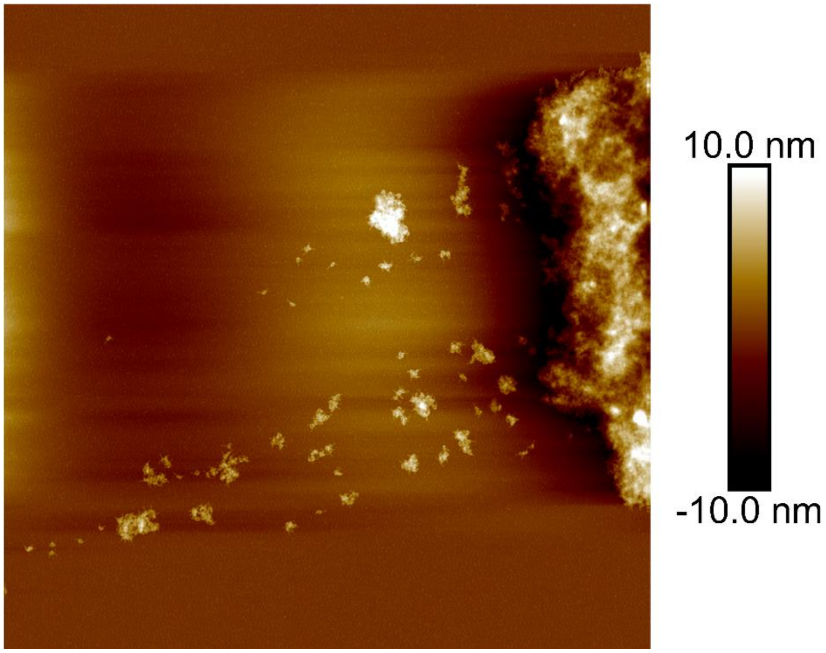


**Figure S10**: AFM image of aggregation of the NuSpring DNA Origami structure folded in the presence of PEG. The inset shows well folded structures. Scalebar: 2 µm.

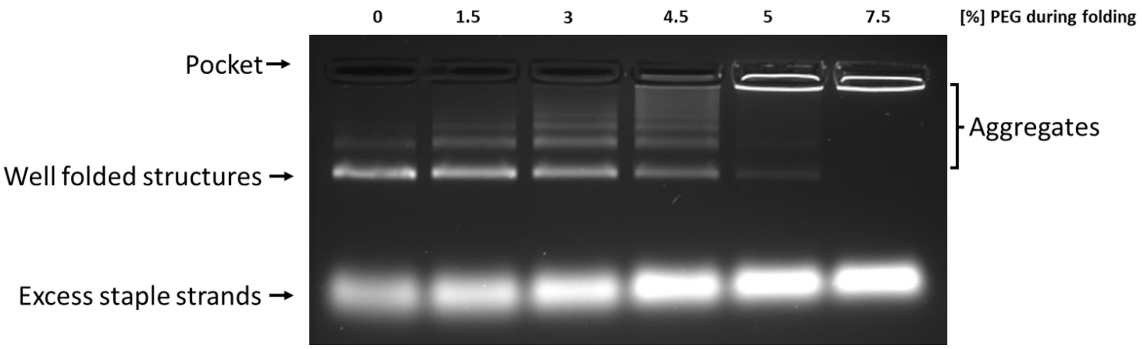


**Figure S11**: Agarose gel electrophoretic analysis of the NuSpring DON folded using a standard folding protocol but with different amounts of PEG added. The aggregation at higher PEG percentage indicates that the PEG present in purification supernatant makes folding with staples directly from this supernatant unfeasible.

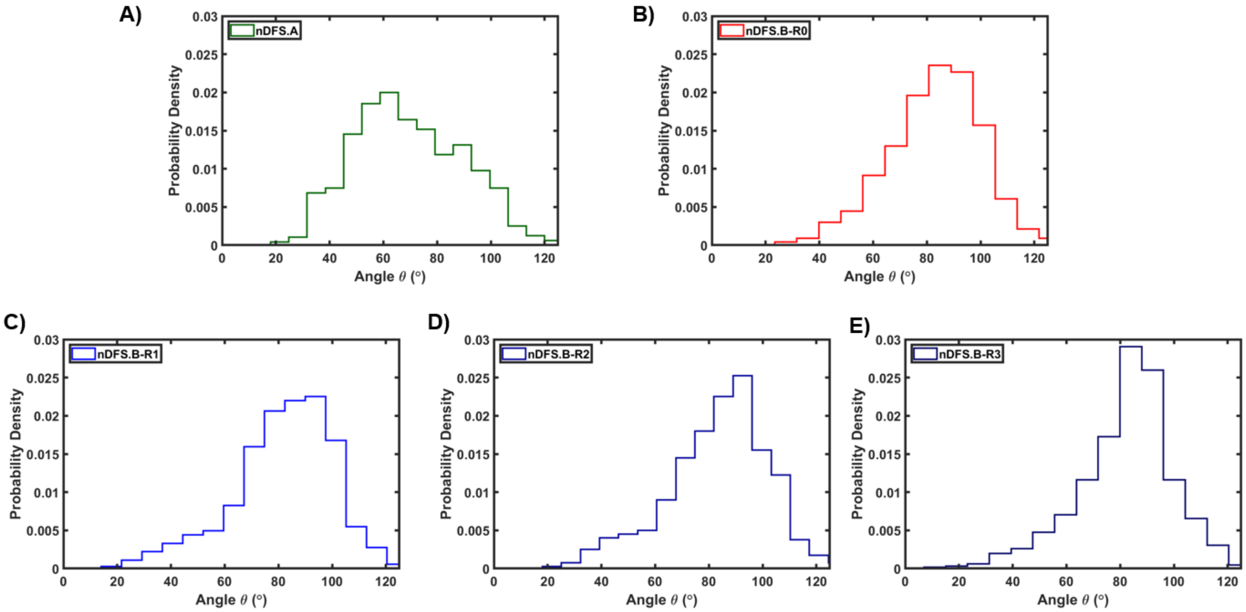


**Figure S12:** Angular distributions of nDFS.A, nDFS.B, and recycled nDFS.B DONs measured from TEM images. A) through C) comprise the data shown in Figure 3 C), while in D) and E) the structures were folded with further recycled staples up to 3 rounds.

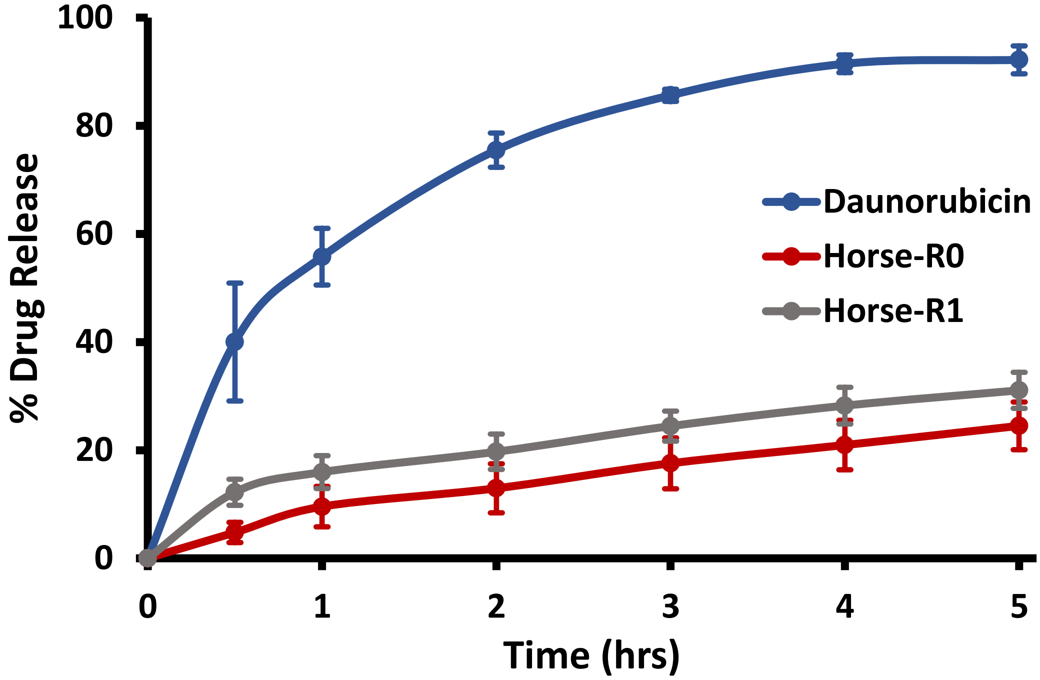


**Figure S13:** Drug release profile using pristine and recycled Horse DONs. Absorbance was measured at 30 minutes and 1-hour time points up to 5 hours, and drug release was calculated by comparing absorbance at time point to absorbance at initial time point. The experiments were performed in triplicate.

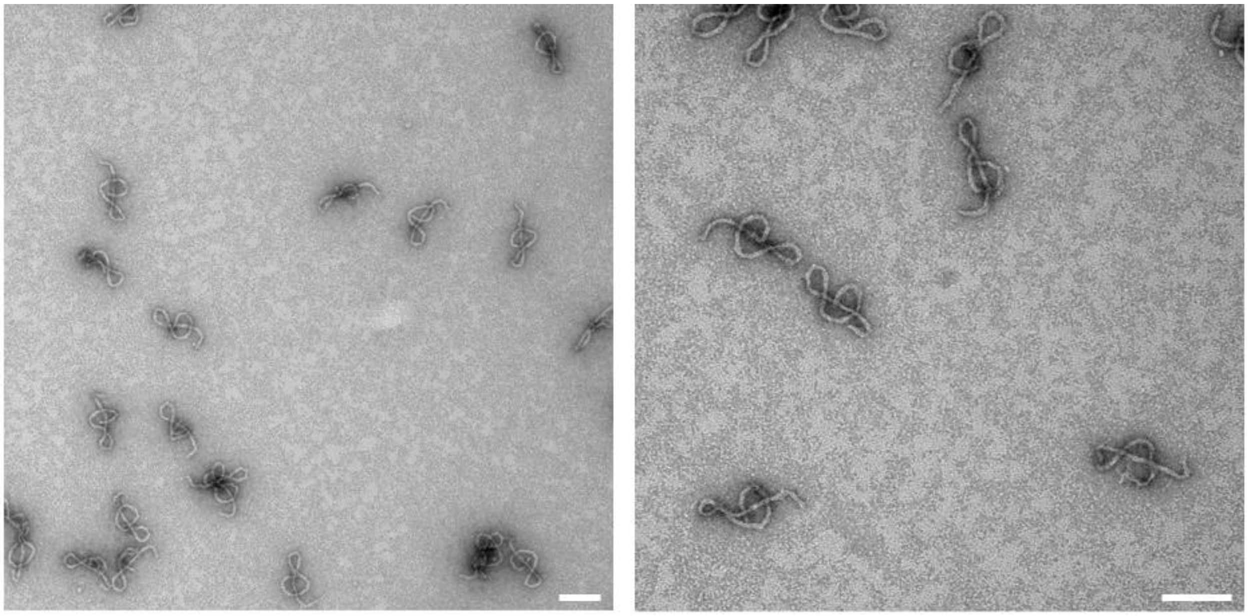

**Figure S14**: Uncropped TEM images of minimal waste fabrication G Clef structures (reprogrammed from NuSpring with recovered staples). Scalebars: 100 nm.

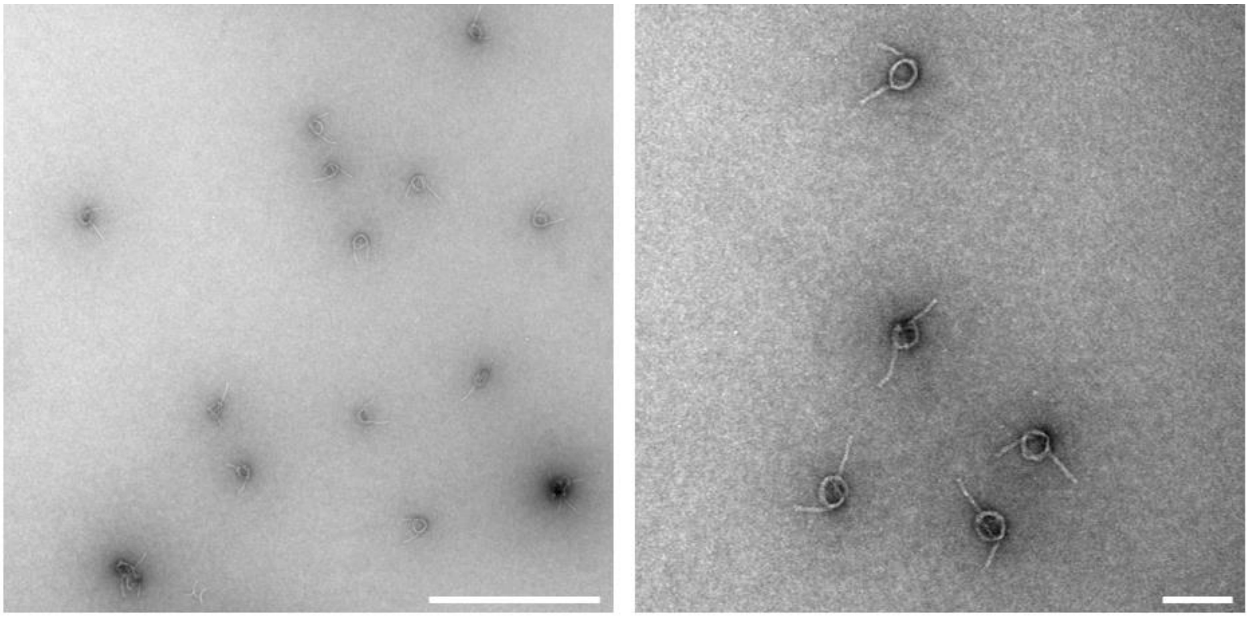

**Figure S15**: Uncropped TEM images of minimal waste fabrication NuSpring structures (reprogrammed from G-Clef with recovered staples). Scalebars: 500 nm (left) and 100 nm (right).

**Supplementary Tables**

| Structure | Yield [%] |
|---|---|
| N | 88 |
| GN | 86 |
| H | 97 |
| GNH | 89 |
| G | 93 |
| GNHG | 81 |

**Table S1:** Yield of cyclically reprogrammed structures by gel fraction folded. Yield was calculated from the gel in Figure S1. To obtain the yield, the main structure band intensity was integrated and divided by the total intensity from the lane integrated from the structure band up to the loading well. This gel analysis was done using GelAnalyzer 23.1.1 (GelAnalyzer 23.1.1 by Istvan Lazar Jr., PhD and Istvan Lazar Sr., PhD, CSc) and baseline detection was performed using a Peak width tolerance of 70.

| Structure | Yield [%] |
|---|---|
| R0 | 91 |
| R1 | 90 |
| R2 | 91 |
| R3 | 90 |

**Table S2:** Yield of structures recycled by staple recovery over several rounds calculated by gel fraction folded Yield was calculated from the gel in Figure S1. The process for calculating the yield is described above in **Table S1.**

| Complementary Length (bp) | nDFS.B-R0 | nDFS.B-R1 |
|---|---|---|
| 1 | -0.020 +/- 0.001 | -0.017 +/- 0.001 |
| 5 | 0.125 +/- 0.004 | 0.122 +/- 0.002 |
| 6 | 0.288 +/- 0.002 | 0.265 +/- 0.002 |
| 7 | 0.509 +/- 0.002 | 0.465 +/- 0.005 |
| 8 | 0.577 +/- 0.006 | 0.529 +/- 0.002 |
| 9 | 0.630 +/- 0.006 | 0.588 +/- 0.002 |
| 10 | 0.673 +/- 0.003 | 0.630 +/- 0.001 |
| 14 | 0.836 +/- 0.001 | 0.805 +/- 0.005 |

**Table S3:** FRET values of pristine and recycled nDFS.B samples with fluorescent latch described in Figure S8B. Each sample was measured three times with the mean and standard deviation reported above. FRET was calculated using the RatioA method described by Clegg et. al (6).

**Table S4: Standard Thermal annealing protocol (TI)**

| T [°C] | min / °C |
|---|---|
| 65 | 15 |
| 64-61 | 3 |
| 60 | 5 |
| 59-58 | 10 |
| 57 | 15 |
| 56 | 25 |
| 55 | 30 |
| 54 | 45 |
| 53-49 | 60 |
| 48-45 | 42 |
| 44 | 36 |
| 43-42 | 32 |
| 41-39 | 20 |
| 38 | 15 |
| 37 | 10 |
| 36-35 | 5 |
| 34-30 | 2 |
| 20 | stay |

**Table S5: Reprogramming Thermal annealing protocol (TR)**

| T [°C] | min / °C |
|---|---|
| 95 | 1 |
| 65 | 15 |
| 64-61 | 3 |
| 60 | 5 |
| 59-58 | 10 |
| 57 | 15 |
| 56 | 25 |
| 55 | 30 |
| 54 | 45 |
| 53-49 | 60 |
| 48-45 | 42 |
| 44 | 36 |
| 43-42 | 32 |
| 41-39 | 20 |
| 38 | 15 |
| 37 | 10 |
| 36-35 | 5 |
| 34-30 | 2 |
| 20 | stay |
